# CB1 receptor inhibition in fragile X syndrome mice impacts alternative splicing alterations in hippocampal synaptoneurosomal transcriptome

**DOI:** 10.1101/2025.07.01.662539

**Authors:** Lucía de los Reyes-Ramírez, Laura Ciaran-Alfano, Marina Reixachs-Solé, Araceli Bergadà-Martínez, Lorena Galera-López, Sara Martínez-Torres, Alba Navarro-Romero, Rafael Maldonado, Eduardo Eyras, Andrés Ozaita

**Affiliations:** Laboratory of Neuropharmacology-NeuroPhar, Department of Medicine and Life Sciences, Universitat Pompeu Fabra, Barcelona, Spain; Research group in Biology of Cognition, Department of Medicine and Life Sciences, Universitat Pompeu Fabra, Barcelona, Spain; EMBL Australia Partner Laboratory Network at the Australian National University, Canberra, Australia; The John Curtin School of Medical Research, Canberra, Australia; Hospital del Mar Medical Research Institute (IMIM), Barcelona, Spain

**Keywords:** Fragile X syndrome, synaptoneurosomes, CB1 receptor, splicing

## Abstract

**Background:** Fragile X syndrome (FXS) conveys the most frequent heritable genetic cause of intellectual disability and autism. It is caused by a CGG repeat expansion in *FMR1* gene that leads to the loss of fragile X messenger ribonucleoprotein 1 (FMRP). FMRP is highly abundant in synapses, where regulates mRNAs to maintain synaptic plasticity. Treatments under development significantly ameliorate neurological and behavioral landmarks in the mouse model of the disorder, the *Fmr1* knockout (FX) mouse. Specifically, previous studies revealed that pharmacological and genetic inhibition of cannabinoid type-1 receptor (CB1R) restored phenotypic traits in FX mice. However, the molecular hallmarks associated with this experimental therapeutic intervention are largely unknown.

**Methods:** First, we aimed to evaluate the validity of synaptoneurosomes preparations to investigate specific mRNA modifications at synapses. Afterwards, combining *in silico* high-throughput analysis and biochemical determinations, we analyzed the hippocampal synaptoneurosomal transcriptome after pharmacological inhibition of CB1R with the specific antagonist/inverse agonist rimonabant in FX male mice

**Results:** We verified that synaptoneurosomes provide an accurate representation of synaptic composition and function. Then, we found that rimonabant treatment had a limited impact at gene expression level but produced significant modifications in transcript expression. Indeed, detailed analysis of alternative splicing events revealed a relevant number of events in which splicing was reverted from the FX form to the WT form by the treatment.

**Limitations:** We demonstrated that rimonabant treatment alters the AS landscape in FX hippocampal synaptoneurosomes; however, further studies are needed to elucidate if other neural components also contribute to the modifications and whether the findings are specific to rimonabant treatment or to CB1R inhibition at synapses. In addition, additional research could be required to clarify whether the changes in AS events could be affected by interindividual variability or technical protocols.

**Conclusions:** We determined that the AS landscape is modified in FX hippocampal synaptoneurosomes and that these changes are sensitive to rimonabant treatment which could explain the beneficial effects of this experimental therapeutic approach in FXS. Altogether, our results reveal a new level of complexity in the effect of pharmacological treatment to improve symptoms in the context of FXS.

## Background

Fragile X syndrome (FXS) is the most prevalent inherited cause of intellectual disability and the most common monogenic cause of autism spectrum disorder (ASD) with an incidence of 1:4,000 males and 1:8,000 females [1]. It is caused by a trinucleotide CGG expansion in the 5’ untranslated region of the X-linked fragile X messenger ribonucleoprotein 1 gene (*FMR1*), which leads to *FMR1* hypermethylation and the subsequent loss of its related protein, FMRP [2]. FMRP is an RNA-binding protein with a key role in local mRNA modulation in both pre- and postsynaptic compartments to maintain synaptic plasticity and connectivity [3]. FMRP also controls the activity of transcription factors, chromatin-modifying enzymes or RNA binding proteins key in RNA production and function [4].

The *Fmr1* knockout (FX) mouse model [5] is the most widely used mouse model to study FXS. This model recapitulates some phenotypic traits of subjects with FXS, such as mild cognitive deficits, hyperactivity, or autistic-like behavior [6]. It also reproduces neuronal abnormalities including the alterations in density and morphology of dendritic spines [7] or the elevated group I mGluR5-dependent long-term depression (LTD) in the hippocampal CA1 region [8]. In addition, this model allowed describing the role FMRP plays in the regulation of RNA alternative splicing (AS) in different brain regions and peripheral tissues (Shah et al., 2020; Jung et al., 2023). Notably, alterations in AS are a major source of pathological changes in psychiatric and neurological diseases, including autism spectrum disorder [11], although the specific alterations in FXS are largely unknown.

The endocannabinoid system (ECS) is an endogenous neuromodulatory network with a relevant role in cognition-associated processes such as synaptic plasticity (Zou & Kumar, 2018). Specifically, cannabinoid type-1 receptor (CB1R) can be pharmacologically or genetically inhibited to improve memory performance in mouse models of FXS [14,15]. Indeed, it was previously described that CB1R pharmacological inhibition with a very low dose of the specific systemic CB1R antagonist/inverse agonist rimonabant or the specific CB1R neutral antagonist NESS0327; or the genetic inhibition of CB1R, ameliorated cognitive deficits in FX mice [14,15]. In addition, low doses of rimonabant resolved the aberrant mGluR5-LTD, the enhanced mammalian target of rapamycin signaling and the altered hippocampal spine density in FX mice [14,15]. Nevertheless, the molecular hallmarks underlying the behavioral and neurological improvements of rimonabant treatment in FX mice remain unknown.

In the present study, we demonstrate that the use of hippocampal synaptoneurosomes allows performing a thorough assessment of synaptically relevant mRNA species. Furthermore, we reveal that in the hippocampal synaptic transcriptome of FX mice there is a complex modulation at transcript level. Interestingly, analysis of AS events revealed a significant set of changes in FX samples that were sensitive to CB1R inhibition with a very low dose of rimonabant. Altogether, the present study uncovers a previously unknown level of complexity in the effects of treatments that improve cognitive performance in FXS.

## Methods

### Ethics

All animal procedures were performed following “Animals in Research: Reporting Experiments” (ARRIVE) guidelines in accordance with standard ethical guidelines [16,17] (European Directive 2010/63/EU) and were approved by the local ethical committee (Comitè Ètic d’Experimentació Animal-Parc de Recerca Biomèdica de Barcelona, CEEA-PRBB).

### Animals

FX mice in Friend Virus B (FVB.129) background (*Fmr1* KO, FVB.129P2-Pde6b+ Tyrc-ch Fmr1tm1Cgr/J) and wild-type mice (WT, FVB.129P2-Pde6b+ Tyrc-ch/AntJ) were purchased from The Jackson Laboratory and crossed to obtain FX and WT littermates used in this study. Young adult male mice (8-15 weeks old) were used for pharmacological approach. Mice were housed four per cage in a temperature (21 ± 1°C) and humidity (55 ± 10%) controlled environment with food and water available *ad libitum*. Pharmacological treatments were performed during the light phase of a 12 h cycle (lights on at 8 am; light off at 8 pm).

### Drugs

Rimonabant (0.1 mg/kg) from Axon Medchem was diluted in 5% ethanol, 5% cremophor-EL and 90% saline. Rimonabant and its vehicle were injected intraperitoneally (i.p.) during 7 d in a volume of 10 ml/kg of body weight.

### Hippocampal bulk and synaptoneurosomal preparation from mouse brain

Hippocampal tissues were rapidly dissected on ice, frozen on dry ice and stored at -80°C until used. Frozen hippocampal tissues were Dounce-homogenized by 10 strokes with a loose and 10 strokes with a tight pestle in 30 volumes of synaptoneurosome lysis buffer (in mM) (CaCl2 2.5, NaCl 124, KCl 3.2, K2H2PO4 1.06, NaHCO3 26, MgCl2 1.3, glucose 10, sacarose 0.32, HEPES/NaOH 20, pH 7.4) including phosphatase inhibitors (NaPyro 5, NaF 100, NaOrth 1, β-glycerolphospate 40) and protease inhibitors (leupeptin 1 mg/ml, aproptinin 10 μg/ml, pepstatin 1 μg/ml, phenylmethylsulfonyl fluoride 1mM). Then, 35 μl of volume were collected and stored at -80°C as bulk hippocampal crude homogenate. The remaining volume was processed following synaptoneurosomal enrichment preparation as previously described [18,19]. Briefly, the bulk homogenate was centrifuged at 2,000×g for 1 min to recover the supernatant (S1) and the pellet was resuspended in 1 ml of synaptoneurosomes lysis buffer to further centrifuge at 2,000×g for 1 min. A second supernatant (S2) was recovered and combined with previous one (S1), and the combined supernatant (S1+S2) was filtered through a hydrophobic PTFE 10 μm filter (LCWP02500, Merck Millipore). The filtrate was centrifuged at 4,000×g for 1 min to attain the supernatant (S3) and then centrifuged at 14,000×g for 5 min, to obtain the pellet as the synaptoneurosomes.

### Synaptoneurosomal enrichment preparation

Mouse hippocampal tissues were obtained 24 h after the last rimonabant (0.1 mg/kg, 7 d, i.p.) or vehicle administration and frozen at -80°C. Hippocampal tissues from all groups were processed in parallel to minimize variability following the protocol for synaptoneurosomal fractions described above.

### RNA extraction

Hippocampal bulk and synaptoneurosome RNA were isolated using RNAeasy miniKit (74104, QIAGEN) following manufacturer’s instructions.

RNA concentration, integrity and purity were measured using NanoDrop (Thermo Fisher Scientific) and Agilent 2100 Bioanalyzer (Agilent Technologies). RNA concentration and quality was uniform between samples and met the criteria of RIN > 6.8.

### RNA Sequencing analysis

RNA libraries from hippocampal synaptoneurosomes and bulk hippocampus samples generated from the same animal were obtained by Macrogen Inc. using SMARTer Ultra Low Input kit (Takara Bio/Clontech, 634940) according to manufacturer’s instructions and samples were sequenced using Illumina NovaSeq6000 flow cell. WT and FX hippocampal synaptoneurosomal samples were processed at the Universitat Pompeu Fabra Genomics Facility. RNA libraries were generated using NEBnext Ultra Directional RNA library prep kit for Illumina (New Englands Biolabs) following the manufacturer’s protocols in combination with Poly(A) mRNA magnetic isolation module (NEB) and were sequenced using Illumina NextSeq500 high-throughput flow cell. All raw RNA-seq data were deposited on the Gene Expression Omnibus (GEO) database and made publicly available with the identifier GSE284627 for hippocampal synaptoneurosomes and bulk hippocampus and GSE285032 for WT and FX hippocampal synaptoneurosomes.

Pseudogenes were removed from Ensembl GRCm39 annotation to further obtain transcript abundance in transcripts per million (TPM) using *Salmon* 1.5.2 [20] in its quasi-mapping-based mode. Gene and transcript level quantification in counts per million (CPM) were obtained using *tximport* (v1.18) R package [21].

### Differential expression analysis

Low-level expressed genes and transcripts were filtered out based on mean CPM > 1. Differential expression analyses at gene and transcript level were performed using *DESeq2* (v1.30) Bioconductor package [22]. Four comparisons were made: synaptoneurosome transcriptome (bulk hippocampus vs. synaptoneurosomes, HC-BULK *vs.* HC-SYN), genotype effect (WT-VEH *vs.* FX-VEH), treatment effect in FX mice (FX-VEH *vs.* FX-RIM) and rimonabant effect in WT mice (WT-VEH *vs.* WT-RIM).

Differential expression was estimated based on fold-changes, and p-values were corrected using the Wald test procedure, with a significance cut-off set as |log_2_FC| > 0.2 and adjusted *p* < 0.1 for differential gene expression (DGE) analysis and adjusted *p* < 0.05 for differential transcript expression (DTE) analysis. Overrepresentation enrichment analyses were performed to investigate synaptoneurosomes composition with the synapses specific database *SynGO* [23] against brain expressed background, setting medium stringency and top and second level terms as labels for cellular component analysis. To analyze the function of the genes associated with differentially expressed transcripts, enrichment analysis was performed with *clusterProfiler* (v3.14) Bioconductor package [24]. Each differentially expressed gene set was analyzed separately using all synaptoneurosomes expressed genes as background. Benjamini-Hochberg multiple testing procedure was used to calculate the p-value for the functional annotation, and results were determined statistically significant at an adjusted *p* < 0.05 for each gene ontology (GO) term.

### Alternative splicing analysis

AS analysis was performed using *SUPPA* [25,26] to predict the relative inclusion and differential splicing of different types of AS events (Supplementary fig. 1A). The relative inclusion of each event for all experimental groups was calculated as Percent Spliced In (PSI) using the TPMs previously obtained. To test the differential inclusion in each comparison, delta PSI (dPSI) was calculated as the difference of the mean PSI from each group: genotype effect (PSI_FX-VEH_ – PSI_WT-VEH_), treatment effect in FX (PSI_FX-RIM_ – PSI_FX-VEH_) and rimonabant effect in WT (PSI_WT-RIM_ – PSI_WT-VEH_). The significance of each event was calculated using a linear regression model and p-values were corrected by calculating the false discovery rate (*FDR*). An event was considered significant when |dPSI| > 0.05 and adjusted *FDR* < 0.05.

### Specific assays for alternative splicing events

To detect AS events, we designed custom TaqMan gene expression assays (4331348, Applied Biosystems) for both the inclusion and the exclusion of each AS event (Supplementary fig. 1B). Primers and internal probes for each inclusion/exclusion event were designed using the Primer Express (4363991, Applied Biosystems) software. Primer dimerization and specificity were checked with OLIGO [27] and PrimerBLAST [28] tools.

### Gene expression and alternative splicing analysis by RT-PCR

Comparable amounts of RNA from hippocampal synaptoneurosomes samples were reverse transcribed to cDNA using the High-capacity cDNA Reverse Transcription kit (4368814, Applied Biosystems) following the manufacturer’s instructions. The cDNAs were stored at −20°C until use. Quantitative real time PCR (qRT-PCR) was carried out in triplicate with QuantStudio 12K Flex Real-Time PCR System (4471134, Applied Biosystems) using SYBR Green I (04707516001, Roche) or TaqMan chemistry (4369016, Applied Biosystems) PCR Master Mix.

For gene expression, qRT-PCR was performed in 10 μl reaction using 0.1 ng/μl of cDNA with the previously described primers:

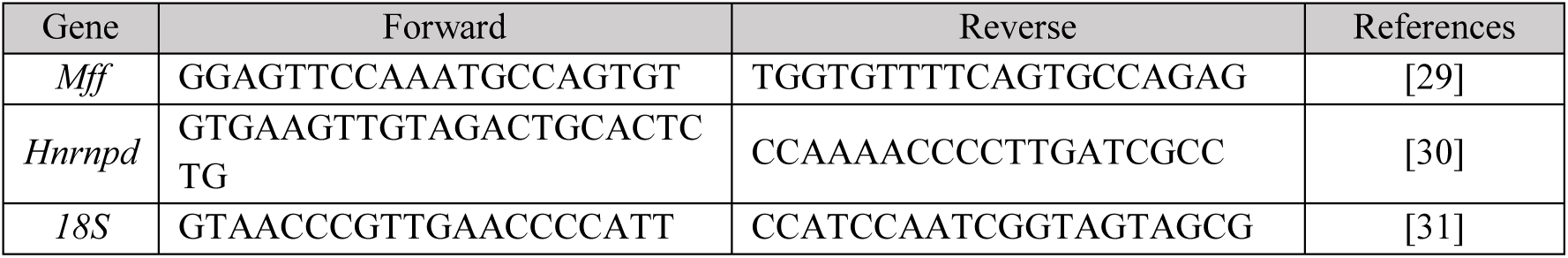

To determine splicing event expression, qRT-PCR was performed in 5 μl reaction using 0.08 ng/μl of hippocampal cDNA. Customized TaqMan probes (Applied Biosystems) were used to detect each splicing event and probe for *18S* RNA (Hs99999901_s1) was used as endogenous control for hippocampal normalization. Quantification was performed using the comparative double-delta Ct method [32] and the fold change was calculated using the equation 2^-ΔΔCt^.

To estimate the relative gene and inclusion/exclusion expression, the control sample (WT-VEH) was standardized at a value of 1. To estimate each splicing events expression, an experimental differential Percent Spliced In (experimental dPSI) was calculated from inclusion and exclusion expression values, similar to the *in silico* calculation: inclusion expression levels from each event were divided by the sum of inclusion and exclusion expression levels for that event.

### Statistical analysis

Data was analyzed with RStudio (v4.0) and GraphPad Prism (v8.0) software. Molecular results were reported as mean ± standard error of the mean (s.e.m). Statistical comparisons were evaluated using unpaired Student’s t-test for two groups comparisons or one-way ANOVA for multiple comparisons. Subsequent Tukey analysis was used when required (significant interaction between factors). Comparisons were considered statistically significant when *p* < 0.05.

## Results

### Synaptoneurosomes exhibit robust representation of the synaptic transcriptome

We used high-throughput RNA sequencing to identify gene populations in hippocampal synaptoneurosomes (HC-SYN) compared to bulk hippocampus (HC-BULK) to assess whether the synaptoneurosomal preparation improves the detection of synaptically-relevant RNA species. Both types of RNA samples (HC-SYN and HC-BULK) were obtained from three mice starting from a single piece of hippocampal tissue that was split during sample processing to assess relative gene enrichment. We found a relevant effect of subcellular fractionation when comparing sample-to-sample variation in gene expression (Fig. 1A), pointing to the type of subcellular fractionation (bulk hippocampus *vs.* synaptoneurosomes) instead of mouse of origin, as the main characteristic influencing transcriptome identity. DGE analysis comparing synaptoneurosomes and bulk hippocampus (HC-BULK *vs.* HC-SYN), revealed 749 genes enriched in synaptoneurosomes while 1,019 genes were depleted in the same samples (Fig. 1B). Gene enrichment analysis showed that among the top-fifteen enriched GO cellular component terms, most of them were related to synapses and transmembrane transport (Fig. 1C). Moreover, among all enriched biological process GO terms, more than half of them (54%, 154/287) were associated with synaptic organization, synaptic plasticity, and mitochondrial activity, giving consistency to our strategy to study the synaptic transcriptome. Additionally, we analyzed synaptic processes and locations using the synaptic function annotation database *SynGO*. Specifically, we found that a relevant number of the genes significantly enriched in synaptoneurosome samples compared to bulk hippocampus, were synapse-associated genes (Fig. 1D).

**Figure 1:**
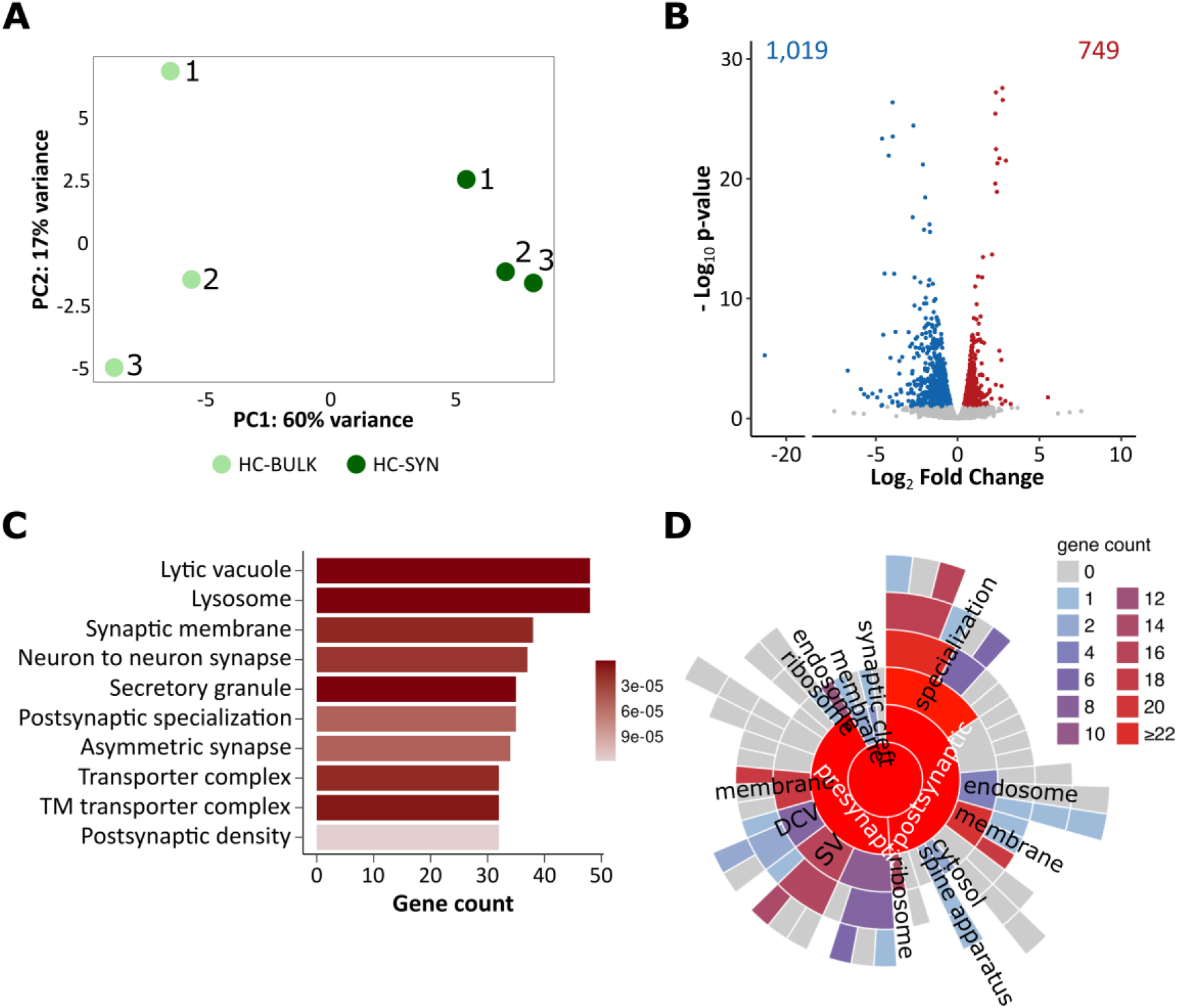
Transcriptomic profile of hippocampal synaptoneurosomes significantly differ from bulk hippocampal transcriptome. **(A)** Principal Component Analysis (PCA) of sequenced samples from the same animal colored based on sample type (HC-SYN, *n* = 3; HC-BULK, *n* = 3). **(B)** Volcano plot of differentially expressed genes from hippocampal synaptoneurosome samples compared to bulk hippocampal samples. (HC-BULK *vs.* HC-SYN). Blue dots represent 1,019 significantly depleted genes (log_2_FC < - 0.2 and adjusted *p-value* < 0.1) and red dots 749 significantly enriched genes (log_2_FC > 0.2 and adjusted *p-value* < 0.1). **(C)** Gene ontology enrichment analysis for enriched genes in hippocampal synaptoneurosomes. GO terms represent only cellular components (CC). **(D)** Sunburst plot of synaptic annotated ontologies for location (CC) terms. Colors are based on gene counts, including child terms. TM, transmembrane.

### Sub-chronic CB1R inhibition with rimonabant shows the reversion in the expression of Mff, Hnrnpd and Slain2 in FX synaptoneurosomes

To determine molecular changes at synaptic contacts associated with the lack of FMRP and the effect that CB1R inhibition could have, we performed high-throughput RNA sequencing in synaptoneurosomes obtained from hippocampus. By comparing FX and WT mice treated with vehicle (FX-VEH and WT-VEH, respectively), we found 29 genes upregulated, and 19 genes downregulated (Fig. 2A). Out of these genes, we observed the upregulation of genes such as *Pcdhgb5* or *Gpr37* that were associated with dendritic arborization [33] and ERK phosphorylation cascade [34], respectively; and the downregulation of *Fkbp5* which has an effect in synaptic transmission [35] or *Rheb* that is implicated in mTORC1 signaling [36] but these set of genes did not produce any hit when gene enrichment analysis was performed, pointing to limited impact of the *Fmr1* KO mutation in the synaptic transcriptome in terms of gene expression.

**Figure 2:**
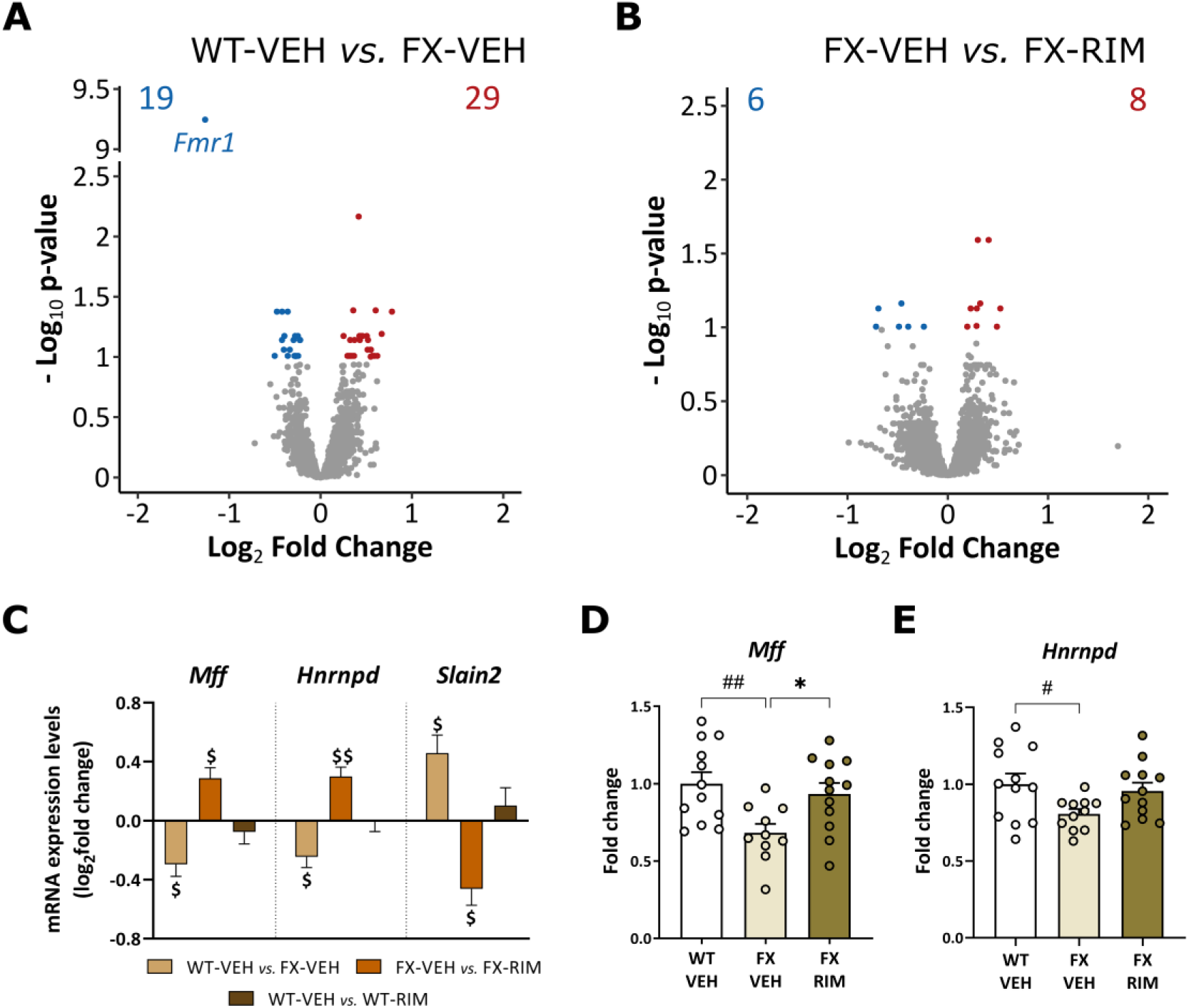
Systemic CB1R inhibition modulates the expression of *Mff* and *Hnrnpd* in FX hippocampal synaptoneurosomes. Volcano plot of differentially expressed genes in **(A)** FX synaptoneurosomes compared with WT (WT-VEH vs. FX-VEH) and in **(B)** FX synaptoneurosomes treated with rimonabant (0.1 mg/kg, 7 d) or vehicle (FX-VEH vs. FX-RIM). Red and blue dots represent significantly up or downregulated genes, respectively. (WT-VEH, n = 4; FX-VEH, n = 5; FX-RIM, n = 6). Cut-off was set at |log2FC| > 0.2 and adjusted *p*-value < 0.1. **(C)** Expression levels of differentially expressed genes that revert their expression after rimonabant (0.1 mg/kg, 7 d) (RIM) administration in FX mice. Data obtained from next-generation RNA-seq are expressed as fold change ± s.e.m $ *p* < 0.1, $$ *p* < 0.05 (in each comparison) by Benjamini-Hochberg adjustment following Wald test analysis. **(D,E)** Change in relative mRNA level from hippocampal synaptoneurosomes measured by qRT-PCR for **(D)** *Mff* (*n* = 10-12) and **(E)** *Hnrnpd* (*n* = 11-12). Data are expressed as mean ± s.e.m. # *p* < 0.05, ## *p* < 0.01 (genotype effect); * *p* < 0.05 (treatment effect) by one-way analysis of variance (ANOVA) with Tukey’s *post hoc* test.

Then, we analyzed the effect of sub-chronic systemic CB1R inhibition with rimonabant in FX and WT synaptoneurosomes. Rimonabant (0.1 mg/kg, 7 d) treatment modified the expression of 14 genes in FX synaptoneurosomes (Fig. 2B). Notably, in WT synaptoneurosomes, rimonabant uniquely upregulated *Col3a1* and *Cplan1* and downregulated *Pde2a*, none of them in common with the effect of rimonabant in FX samples (Supplementary fig. 2A).

Next, we overlapped the gene expression profiles from the previous comparisons to determine the specific effect of rimonabant in FX mice. With this overlap analysis, we identified three genes that were present in both comparisons with an opposite expression: *Mff*, *Hnrnpd* and *Slain2* (Fig. 2C). To further validate this reversion produced by rimonabant in FX, we assessed *Mff* and *Hnrnpd* mRNA expression levels in a different set of samples after rimonabant (0.1 mg/kg, 7 d) administration. qRT-PCR analysis confirmed a decrease in the expression of *Mff* in FX mice that was reverted after sub-chronic treatment with rimonabant (one-way ANOVA, interaction: F (2,31) = 5.58, *p* = 0.008; *post hoc* Tukey, WT-VEH *vs.* FX-VEH *p* = 0.008; FX-VEH *vs.* FX-RIM *p* = 0.04) (Fig. 2D), and a decrease in the expression of *Hnrnpd* mRNA levels in FX mice (one-way ANOVA, interaction: F (2,32) = 3.41, *p* = 0.04; *post hoc* Tukey, WT-VEH *vs.* FX-VEH *p* = 0.04) (Fig. 2E).

### Differential transcript expression analysis reveals a significant effect of CB1R inhibition in FX mice

Based on the results of *Hnrnpd* in DGE and its described function as a direct mRNA regulator [37], we performed the DTE analysis with the same transcriptomic data sets as above.

FX mice showed 254 differentially expressed transcripts compared with WT mice: 145 transcripts were upregulated in FX while 109 were downregulated (Fig. 3A). Gene enrichment analysis of differentially expressed transcripts revealed an over-representation of synapse structure and synapse organization genes in both up and downregulated transcripts (Fig. 3B-C). Among the downregulated transcripts, we also observed an enrichment in biological processes associated with mRNA processing, such as regulation of mRNA metabolic process or RNA splicing (Figure 3B). Notably, within the 109 transcripts downregulated in FX synaptoneurosomes, we found some splicing factors such as *Celf2* or members of *Hnrnp* family, pointing to a reduction in transcripts needed for mRNA processing and transcripts that may undergo splicing [38].

**Figure 3:**
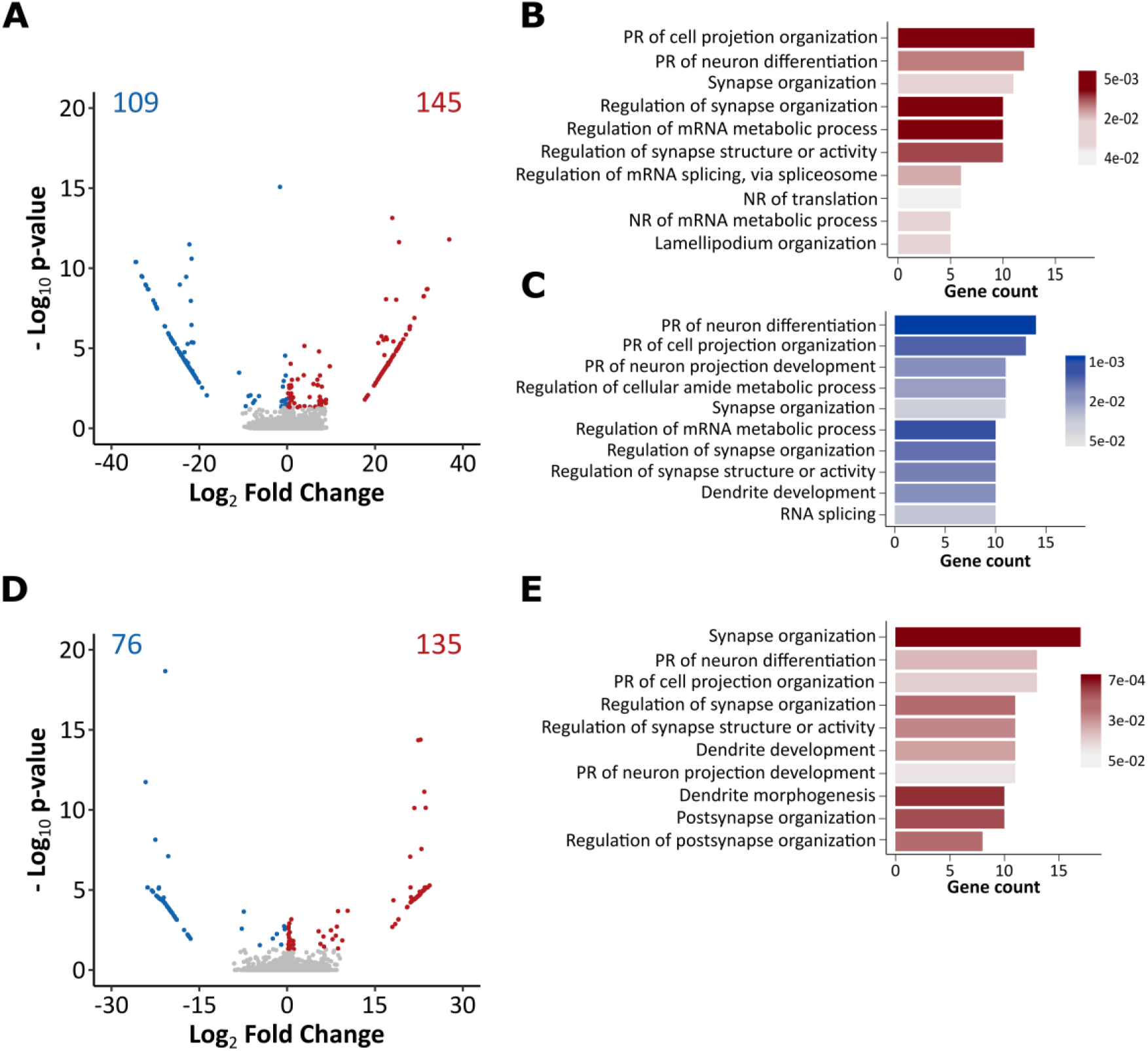
Transcriptomic landscape of FX hippocampal synaptoneurosomes at transcript level after vehicle or rimonabant administration. **(A)** Volcano plot of differentially expressed transcripts in **(A)** FX compared to WT (WT-VEH *vs.* FX-VEH). Red and blue dots represent significantly up or downregulated genes, respectively. **(B,C)** Bar plot showing the top-ten most representative GO terms associated with **(B)** up and **(C)** downregulated transcripts in FX synaptoneurosomes. GO terms represent only biological processes (BP). **(D)** Volcano plot of differentially expressed transcripts in FX synaptoneurosomes treated with rimonabant (0.1 mg/kg, 7 d) (RIM) compared with those treated with vehicle (FX-VEH *vs.* FX-RIM). **(E)** Bar plot showing the top-ten most representative GO terms from biological processes associated with upregulated transcripts in FX treated mice. For volcanos, cut-off was set at |log_2_FC| > 0.2 and adjusted *p-value* < 0.05 (WT-VEH, *n* = 4; FX-VEH, *n* = 5; FX-RIM, *n* = 6). For GO enrichment plots, terms were ordered by the number of genes related with the process and the gradient color is proportional to the significance.

We next analyzed the effect of rimonabant in FX mice at the transcript level. DTE comparing synaptoneurosomes from FX mice treated with rimonabant or vehicle showed 211 transcripts differentially expressed (Fig. 3D); specifically, 135 transcripts were upregulated while 76 were downregulated. Enrichment analysis of the associated genes only showed results among upregulated transcripts, which were related with synapse organization, mRNA splicing and neuron differentiation (Fig. 3E). Interestingly, we found some transcripts from genes such as *Samd4b*, *Acin1* or *Hnrnp* family (*Hnrnpa2b1*, *Hnrnpk* and *Hnrnpd*) associated with mRNA metabolic process and spliceosome function.

We also assessed whether rimonabant administration had similar effects in WT mice. We observed 224 transcripts differentially expressed when we performed the DTE analysis (Supplementary fig. 2B). Gene enrichment analysis showed no functional associations between genes in terms of biological processes and few hits on molecular function and cellular component domains (Supplementary fig. 2C), all very different from those functions observed in rimonabant treated FX mice. This data further indicated that the effects of rimonabant at the transcriptomic level were genotype dependent.

As we did above with gene expression data, we assessed the specific effect of systemic CB1R inhibition in FX synaptoneurosomes by overlapping differentially expressed transcripts from both comparisons (WT-VEH *vs.* FX-VEH and FX-VEH *vs.* FX-RIM). Out of all transcripts differentially expressed in FX synaptoneurosomes, 54% of them changed their expression in the opposite direction after rimonabant treatment (59+27 out of 254). Specifically, 59 downregulated transcripts in FX synaptoneurosomes, were upregulated after rimonabant administration (hypergeometric test, *p* = 1.8e-10) (Fig. 4A); while 27 transcripts were downregulated in the opposite direction (Fig. 4B). Interestingly, while the 27 transcripts that were downregulated in FX mice after CB1R inhibition were not enriched in any gene ontology term, all the terms obtained in the gene enrichment analysis for the 59 transcripts modulated by rimonabant were related with mRNA processing and splicing (Fig. 4C). This reinforced the hypothesis that rimonabant treatment could ameliorate the synaptic alterations observed in FX mice by affecting mRNA splicing machinery.

**Figure 4:**
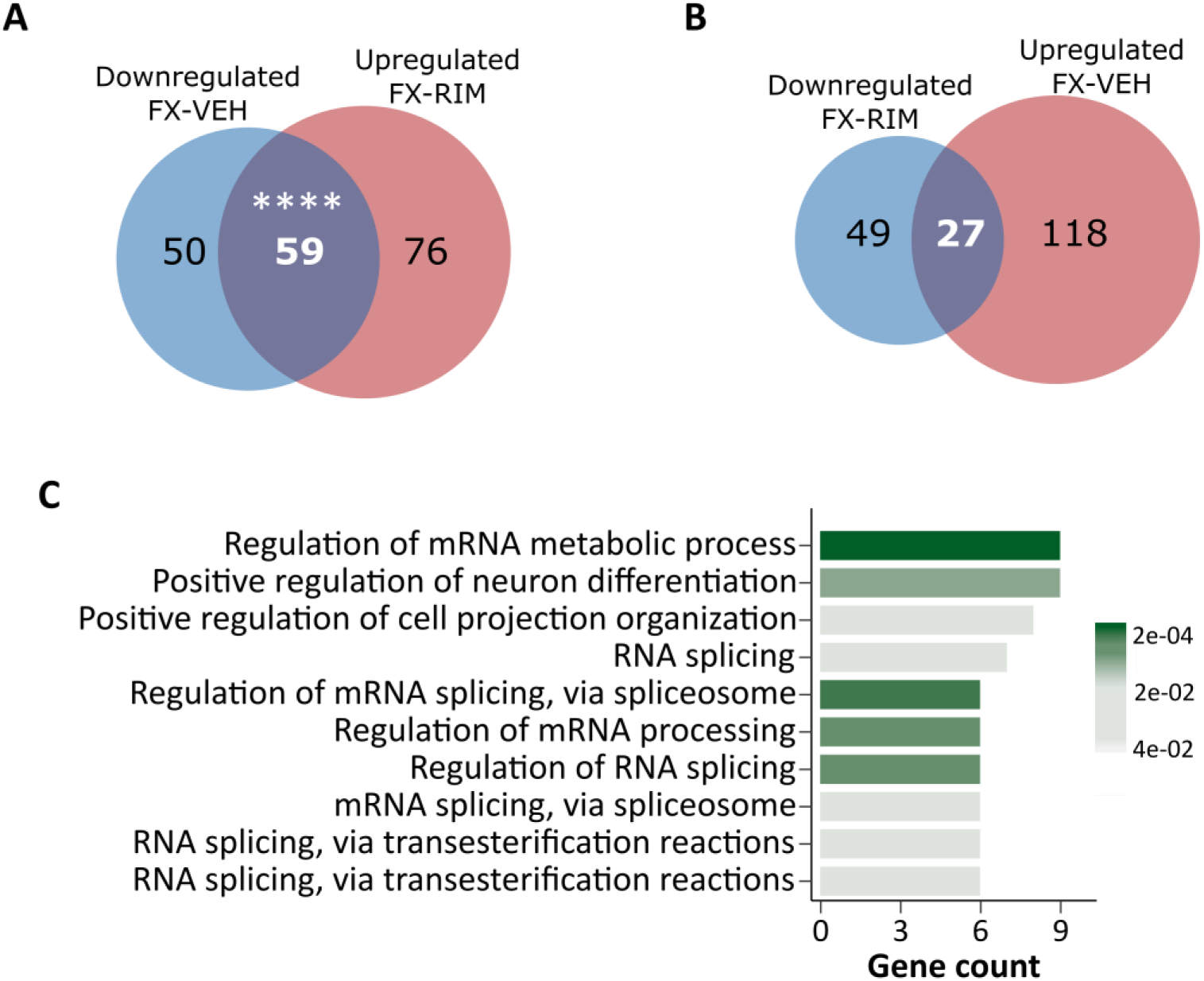
Sub-chronic systemic CB1R inhibition modulates altered transcripts expression in FX and WT hippocampal synaptoneurosomes. Venn diagrams of transcripts that were **(A)** downregulated in FX (WT-VEH *vs.* FX-VEH) and upregulated after rimonabant (0.1 mg/kg, 7 d) treatment (FX-VEH *vs.* FX-RIM) and **(B)** upregulated in FX (WT-VEH *vs.* FX-VEH) and downregulated after rimonabant treatment (FX-VEH *vs.* FX-RIM) **(C)** Bar plot showing the top-ten most representative biological processes associated with genes from downregulated transcripts in FX synaptoneurosomes that were upregulated in FX synaptoneurosomes after rimonabant administration. **** *p* < 0.0001 by hypergeometric test. GO terms were ordered by the number of genes and the gradient color is proportional to the significance.

### FX synaptoneurosomes display a pattern of alternative splicing events that is sensitive to rimonabant administration

Given the implication of the differentially expressed transcripts in mRNA processing together with previous studies associating splicing changes and synaptic specification [11], the possible changes in the AS landscape was investigated in FX synaptoneurosomes. We analyzed AS to obtain direct evidence in terms of skipping exon (SE), mutually exclusive exons (MX), alternative 5’ and 3’ splice-site (A5 and A3), alternative first and last exon (AF and AL) and retained intron (RI) (Supplementary fig. 1A). The lack of *Fmr1* in hippocampal synaptoneurosomes dysregulated the splicing pattern of 206 genes by direct modification of 259 AS events (Fig. 5A, Supplementary fig. 3). AF and SE were by far the most prevalent categories. While only 20% of the genes that we detected to present AS were also described to undergo this type of events in hippocampal slices from FX mice [10], this is the first time that changes in AS were detected specifically in FX synaptoneurosomes. Among the alternatively spliced events between FX and WT mice, we found genes associated with synaptic transmission and synaptic morphology such as *Cpeb2*, *FosB*, *Arf4*, *Cacng5* or *Vdac3*.

**Figure 5:**
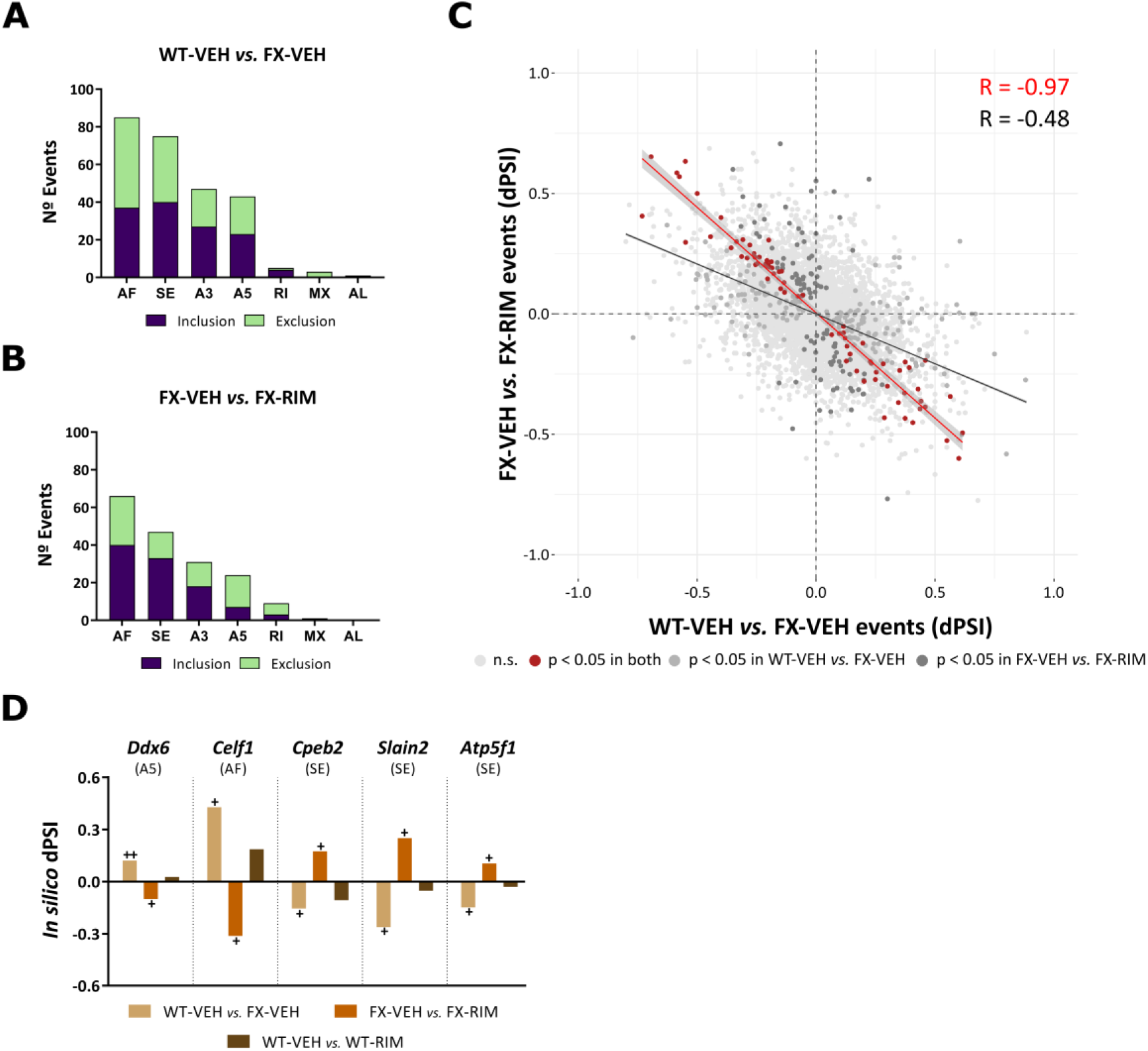
Sub-chronic systemic CB1R inhibition changes the direction of AS events in FX hippocampal synaptoneurosomes. Histogram of AS types identified in **(A)** FX compared with WT and in **(B)** FX treated and non-treated with rimonabant (0.1 mg/kg, 7 d) (RIM). SE, skipping exon; MX, mutually exclusive exons; A5 and A3, alternative 5’and 3’ splice-site, respectively; RI, retained intron; AF and AL, alternative first and last exon, respectively. **(C)** Correlation plot of splicing events identified in both comparisons. In red, significant events in both comparisons (77); in grey and dark grey, significant events only in WT-VEH *vs.* FX-VEH (182) or FX-VEH *vs.* FX-RIM (101) comparisons, respectively. Red line shows the correlation of the red points and grey line the correlation of all other events detected in at least one comparison. Significant events were established based on a |dPSI| > 0.05 and *FDR* < 0.05. **(D)** dPSI values of selected AS events that significantly revert their expression after rimonabant (0.1 mg/kg) (RIM) administration in FX but not in WT mice. Data are expressed as dPSI. + *p* < 0.05, ++ *p* < 0.01 (in each comparison) by linear regression model following *FDR* correction. (WT-VEH, *n* = 4; WT-RIM, *n* = 5; FX-VEH, *n* = 5; FX-RIM, *n* = 6).

We next analyzed AS events in FX synaptoneurosomes treated with rimonabant. This revealed the modification of 178 splicing events in 130 genes compared to non-treated FX synaptoneurosomes (Figure 5B, Supplementary fig. 3). Although fewer events seemed to be regulated by rimonabant, the most representative type of events were again AF and SE. Interestingly, among these events, we found transcripts encoding for ion channels and enzymes that could play a role in gene expression regulation and synaptic transmission such as *Gprin1*, *Kcnip4*, *Gabra5*, *Celf1*, *Mecp2*, *Hnmt* or *Arc*.

To pinpoint the modifications driven by systemic CB1R inhibition, we compared splicing events in synaptoneurosomes from FX-treated mice (FX-VEH *vs.* FX-RIM) with those FX and WT mice treated with vehicle (WT-VEH *vs.* FX-VEH). When we overlapped both sets of AS events, 77 events out of 56 genes were common between genotype and treatment comparisons (Fig. 5C). These events occur in genes associated with RNA processing (*Celf1*, *Ddx6*, *Rbm39*), cytoskeleton modulation (*Slain2, Nek3, Tpm4*), ion channels (*Gprin1, Lrrcd8*), neuronal development (*Psap, Rims3*) and mitochondrial processes (*Atp5pb*) among others. Moreover, we observed a significant negative correlation between the direction and the magnitude of the change in significant events in both FX synaptoneurosomes treated with rimonabant and vehicle (R Pearson = -0.97, *p* < 2.2e-16) (Fig. 5C). Interestingly, the majority of the events that were sensitive to CB1R inhibition in FX mice, were not significantly modified after rimonabant administration in WT mice. Indeed, some interesting events selected for further validation were only significantly detected in FX synaptoneurosomes after vehicle or rimonabant treatment and not in WT synaptoneurosomes after rimonabant administration (Fig. 5D), indicating the specific modulation of these AS events in FXS. Together, we conclude that a significant proportion of the AS events in FX mice could be rescued by systemic CB1R inhibition with a low dose of rimonabant in a genetic context-specific manner.

### Inclusion and exclusion splicing events detected in silico were validated with specific probes

Among the 77 AS events that were modulated in opposite ways between genotype and treatment, we validated *Ddx6, Celf1* and *Cpeb2* using newly generated tissue samples and customized TaqMan assays. As each AS event is described by the exon inclusion and the alternative exon exclusion, we designed specific probes for each transcript according to the type of event (Supplementary fig. 1B). With this experimental design, we measured changes both in the complete event but also in each transcript isoform resulting from AS.

*Ddx6*, which presents an A5 event, encodes DEAD box protein family 6, an RNA helicase associated with translation suppression and mRNA degradation [39]. We found similar results when we measured the expression of the transcripts that defined the exon inclusion and the exon exclusion. *Ddx6* exon inclusion was downregulated in FX and upregulated after rimonabant administration in FX mice (one-way ANOVA, interaction: F (2,33) = 6.46, *p* = 0.004; *post hoc* Tukey, WT-VEH *vs.* FX-VEH *p* = 0.01; FX-VEH *vs.* FX-RIM *p* = 0.006) (Fig. 6A); and the same modulation was observed in *Ddx6* exon exclusion (one-way ANOVA, interaction: F (2,33) = 7.95, *p* = 0.001; *post hoc* Tukey, WT-VEH *vs.* FX-VEH *p* = 0.005; FX-VEH *vs.* FX-RIM *p* = 0.003) (Fig. 6B). In the case of AF event in *Celf1*, an RNA binding protein involved in post-translational modifications [40], transcripts that cover both exon inclusion (one-way ANOVA, interaction: F (2,34) = 8.46, *p* = 0.001; *post hoc* Tukey, WT-VEH *vs.* FX-VEH *p* = 0.03) (Fig. 6C) and exon exclusion (one-way ANOVA, interaction: F (2,33) = 8.05, *p* = 0.001; *post hoc* Tukey, FX-VEH *vs.* FX-RIM *p* = 0.0009) (Fig. 6D) were significantly more abundant in FX mice after rimonabant treatment, although only exon inclusion was also decreased in FX vehicle-treated mice (one-way ANOVA, interaction: F (2,34) = 8.46, *p* = 0.001; *post hoc* Tukey, FX-VEH *vs.* FX-RIM *p* = 0.0007) (Fig. 6C). Lastly, we also tested a SE event described in *Cpeb2*, another RNA binding protein that may regulate synaptic plasticity [41,42]. We found that exon inclusion was less present in FX synaptoneurosomes compared to WT (one-way ANOVA, interaction: F (2,34) = 5.67, *p* = 0.007; *post hoc* Tukey, WT-VEH *vs.* FX-VEH *p* = 0.01) (Fig. 6E) and exon exclusion showed a non-significant tendency to be also less expressed in FX mice (one-way ANOVA, interaction: F (2,33) = 3.23, *p* = 0.051) (Fig. 6F). In addition, exon inclusion was also more present in rimonabant-treated FX mice compared with vehicle-treated FX ones (one-way ANOVA, interaction: F (2,34) = 5.67, *p* = 0.007; *post hoc* Tukey, FX-VEH *vs.* FX-RIM *p* = 0.02) (Fig. 6E). Furthermore, we calculated an experimental dPSI to measure whether the complete event was modified as it was each specific isoform. We found that *Cpeb2* SE event changed in the same direction as *in silico* dPSI: negative in WT-VEH *vs.* FX-VEH and positive in FX-VEH *vs.* FX-RIM (Student’s t-test, *p* = 0.01) (Fig. 6G). However, the other two events did not reach significance. Therefore, we demonstrated that AS changes observed by high-throughput assessment in FX hippocampal synaptoneurosomes, can also be detected in vitro in a different set of samples.

**Figure 6:**
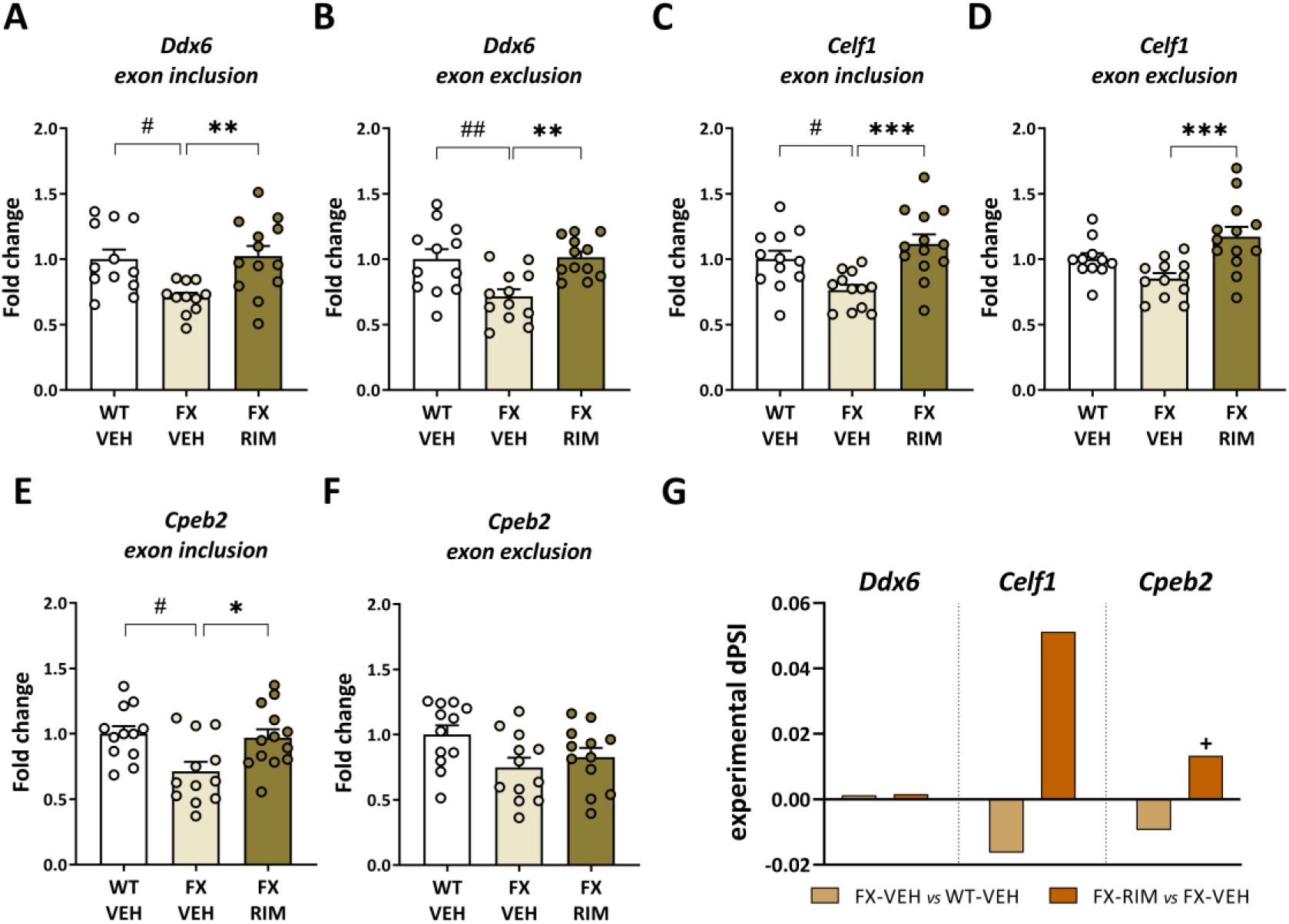
AS event associated-transcripts were modified in FX hippocampal synaptoneurosomes after sub-chronic systemic CB1R inhibition. Hippocampal synaptoneurosomes transcripts that defined (A) *Ddx6* inclusion, (B) *Ddx6* exlcusion, (C) *Celf1* inclusion, (D) *Celf1* exclusion, (E) *Cpeb2* inclusion and (F) *Cpeb2* exclusion were quantified referred to *18S* endogenous control. Data are expressed as mean ± s.e.m. # *p* < 0.05, ## *p* < 0.01 (genotype effect); * *p* < 0.05, ** *p* < 0.01, *** *p* < 0.001 (treatment effect) by one-way analysis of variance (ANOVA) with Tukey’s *post hoc* test. **(G)** Experimental calculation of the differences in PSI (experimental dPSI) in each comparison (WT-VEH *vs.* FX-VEH and FX-VEH *vs.* FX-RIM). Data are expressed as mean of experimental dPSI. + *p* < 0.05 (in each comparison) by Student’s t-test. (*n* = 11-13)

## Discussion

We used RNA sequencing to study the synaptic transcriptome of FX and WT male littermates after 7 d of rimonabant (or vehicle) administration, since this schedule improves cognitive performance and synaptic plasticity in the FXS model [15]. We detected few changes in the overall DGE when WT and FX samples were compared, as well as when comparisons were focused in understanding the effect of rimonabant treatment on FX. Instead, when potential transcripts were independently considered, the analysis revealed a decreased expression of mRNAs involved in synapse organization and RNA processing due to the genetic mutation. Interestingly, detailed study of AS events further demonstrated specific changes associated with the mutation that were reversed by the pharmacological inhibition of CB1R. The present study provides evidence for characteristic RNA splicing events in adult FX hippocampal synapses that are sensitive to CB1R pharmacological inhibition.

Synapses are highly specialized cellular structures that interconnect neurons and allow intercellular communication. The co-isolation of pre and post synaptic structures emerges as a useful technique to understand key molecular mechanisms underlying synaptic plasticity. The solid evidence of mRNA presence and local protein synthesis at synaptic level (Cajigas et al., 2012; Cefaliello et al., 2014; Hafner et al., 2019) allows for both transcriptomic and proteomic approaches to unraveling the molecular mechanisms governing synaptic function. The use of synaptoneurosomes to understand synaptic processes has become particularly relevant for the evaluation of neuronal changes and altered signaling pathways in neurological disorders, including FXS (Reig-Viader et al., 2018; Wang & Savas, 2018). Since FMRP is mainly localized to axons, dendrites and spines acting pre and postsynaptically [47,48], unraveling the impact of its loss at synapses is crucial. We previously characterized our preparation of synaptoneurosomes at proteomic level [18] and now we have validated the use of these synaptoneurosome preparation to obtain a thorough assessment of synaptic mRNA species. One of the major benefits of analyzing synaptic transcriptomes is that low abundant mRNAs are enriched. Indeed, among the enriched genes in hippocampal synaptoneurosomes compared to bulk hippocampus, we found genes linked to synapse architecture, synaptic plasticity, and, interestingly, a large number of genes also connected to mitochondrial bioenergetics. This agrees with a previous study showing that murine synaptoneurosomes were enriched in transcripts associated with synaptic density and ion transport mRNAs, but also energy metabolism- and mitochondria-related mRNAs [49]. Synapses are the cellular compartment with the highest energy consumption [50], as they maintain local protein synthesis, calcium homeostasis, neurotransmitter release and membrane modifications and trafficking [51]. In addition, both processes, synapse organization and mitochondrial function, play essential roles in hippocampal dependent memory [52]. Given that FMRP is able to interact with mRNAs along dendrites and regulates its translation at synapses [4,53], studying mRNA composition in synapses rather than bulk preparations could be relevant to identify synaptic pathways dysregulated in FXS that could be sensitive to pharmacological interventions.

Previous studies in our group have pointed to the inhibition of systemic CB1R to improve memory performance in adult FX mice [14,15]. Specifically, low doses of the systemic CB1R antagonist/inverse agonist, rimonabant, or the neutral CB1R antagonist NESS0327, improved several key features of FX mice including those associated with synaptic structure and plasticity. To understand the synaptic alterations in FX mice as well as the impact of therapeutic intervention directed to CB1R, we analyzed synaptoneurosomes from rimonabant-treated and vehicle-treated FX and WT mice. Notably, gene expression analysis showed few changes between genotype and rimonabant administration, so we paid especial attention to those modifications in which gene expression revealed a change in opposite direction when studying FX synaptoneurosomes after rimonabant or vehicle treatment.

In this regard, we observed a downregulation of *Mff* mRNA expression in FX that was normalized to WT levels with rimonabant treatment. *Mff* product (MFF, mitochondrial fission factor) plays a role in mitochondrial fission, through its interaction with *Drp1* product (DRP1, dynamin related protein 1), regulating mitochondrial size by competition with mitochondrial fusion [54] and therefore controlling not only mitochondrial dynamics but also other functions in the synaptic domain such as neurotransmitter release by limiting Ca^2+^ homeostasis or axonal transport [55]. Notably, changes in mitochondrial fusion/fission mRNA and protein levels were described in FX primary hippocampal neurons, including the reduction in the expression of DRP1 [56], which is consistent with the observed reduction in *Mff* expression. More recently, it has been reported that FMRP regulates MFF translation and that its loss results in mitochondrial fission dysregulation and fragmented mitochondria [57]. Interestingly, it was described how CB1R can regulate mitochondrial dynamics and mitophagy and how CB1R KO mice presented different mitochondria sizes compared to WT mice [58]. These observations are also congruent with our findings on how the dysregulation of *Mff* in FX mice is reversed by rimonabant inhibition of systemic CB1R.

Our results in murine FX synaptoneurosomes also showed downregulation in the expression of *Hnrnpd*. This gene encodes for heterogeneous nuclear ribonucleoprotein D, HNRNP D, also known as AU-binding factor 1, AUF1, one well-characterized RNA binding protein with a suggested role in synapse formation and plasticity [59] through the regulation of the synthesis of several synaptic plasticity-related proteins, early response genes or G protein-coupled receptors [60,61]. Interestingly, the dysfunction of HNRNP proteins was previously associated with neurological diseases, including fragile X tremor ataxia [38,62]. Studies in rat primary neuronal cultures reported an increase in HNRNP proteins at synapses after synaptic stimulus, proposing that these proteins can regulate dendritic spine morphology [63]. Thus, the decrease that we observed in *Hnrnpd* expression could be associated with the elevated hippocampal immature spines observed in FX mice that were significantly reduced after sub-chronic rimonabant treatment [14]. In the absence of FMRP, the excessive and dysregulated mRNA translation could lead to altered synaptic function [64]. This, together with the fact that the reduction in the expression of HNRNP D was also associated with a dysregulation in the stability of mRNAs and subsequent gene expression [65], could suggest a possible relation between FMRP and HNRNP D. Our findings let us hypothesize that the downregulation in *Hnrnpd* expression in FX mice could contribute to the dysregulation in the degradation of the excessive mRNAs that are synthesized in the absence of FMRP.

Nevertheless, gene expression is a complex process in which pre-mRNA splicing and maturation as well as mRNA transport, storage, turnover, and translation, controls the abundance and the location of expressed proteins, and in which HNRNP D has a key role [37]. Based on this, we further analyzed changes in transcript expression in FX hippocampal synaptoneurosomes. Since DTE analysis considers the different versions of transcripts that can be encoded by a single gene, this analysis allows a closer look at the functionality of genes. DTE showed a greater number of hits compared to that at gene level, meaning that in FX mice, changes in RNA biology could critically contribute to the phenotypic alterations and the therapeutic effect of systemic CB1R inhibition. In the FX synaptic transcriptome, we found a dysregulation of transcripts associated with synapse structure and organization in both up and downregulated transcripts. This agrees with previous research in which the lack of FMRP in different tissues has revealed an aberrant expression of relevant genes associated to brain development and structural plasticity (Wang et al., 2023). Moreover, when focusing only in downregulated transcripts, we observed an enrichment in mRNA metabolic processes and RNA splicing. Interestingly, after rimonabant treatment, we observed that transcripts associated with mRNA regulation and spliceosome appeared among the upregulated ones in FX. FMRP is critical in the regulation of RNA metabolism, including AS [67], and some of its targets are RNAs that encode chromatin modifying enzymes [68,69]. These alterations that affect the epigenetic landscape could be associated with an impairment in cognitive function [70], so we hypothesized that the synaptic plasticity normalization observed after systemic CB1R inhibition with rimonabant [14,15] could be associated with changes in the splicing machinery.

We further analyzed the landscape of AS events in all experimental groups and then compared between conditions. Interestingly, we found that only 28 out of 206 genes (∼14%) that undergo splicing in FX synaptoneurosomes are described as FMRP targets according to previous studies in bulk tissues [68,69]. This also supports what we observed at transcript level, that the absence of *Fmr1* expression resulted in the expression reduction of transcripts related to mRNA processing and could be linked to with the changes in AS events. In this regard, many studies show that neurons present a complex cell-type specific program of AS [71,72] and that AS is modulated in subjects with ASD [73,74] and in central and peripheral tissues of FX mice [9,10]. Our data supports that, in the absence of FMRP in hippocampal synaptoneurosomes, mRNA maturation and stability are altered and therefore AS programs that are key in modulating neurogenesis, synapse formation, ion channel activity and synaptic plasticity are also modified [75,76]. Interestingly, some of these splicing changes were sensitive to systemic CB1R inhibition with a low dose of rimonabant. We found a major correlation between the magnitude and the direction of the change between all 77 AS events in FX after rimonabant or vehicle administration. Of note, we found that *Slain2*, a gene involved in axonal growth [77], was alternatively spliced between treated and non-treated FX mice, which could explain the lack of validation at gene expression level. Moreover, the CB1R-sensitive changes in AS of *Atp5pb*, part of ATPase synthase c-subunit, could also be associated with the elevated mitochondrial leak found in FX mice [78] and with the dysregulation of *Mff*.

Notably, most of the genes that were alternatively spliced in FX in a rimonabant-dependent manner were RNA-binding proteins (RBP), which included those that were validated in a different set of samples: *Ddx6*, *Celf1* and *Cpeb2*. We found that the same hippocampal modulations observed by RNA-seq in FX mice, were validated in a different cohort of mice using hippocampal synaptoneurosomes: the detection of *Cpeb2* complete AS event and the modulation of each transcript isoform regarding *Ddx6*, *Celf1* and *Cpeb2*. Interestingly, *Cpeb2* may be linked to memory deficits, since CPEB2 cKO mice exhibit reduced long-term potentiation, defective fear and spatial memory and immature spine morphology [79]. AS changes in *Cpeb2* could be associated with synaptic deficits of FX mice and the fact that rimonabant changed this AS events could be related with the beneficial effect of CB1R in FX mice. Thus, further studies of AS events could help on FXS management and its associated alleviation.

Based on the genes that bear AS events in FX mice, it is unlikely that FMRP is a direct regulator of splicing. Instead, it can mediate RNA processing by regulating other downstream RBP that ultimately regulate key processes for synaptic transmission, such as excitation/inhibition balance [80,81]. This could be in accordance with the described association between alternative isoforms of RBP like *Rbfox1* and changes in this balance [82], a common neurobiological feature of ASD [74,83] and FXS [84]. Given that synapses function and formation can respond to neuronal activity, the AS of these RBP could be somehow associated with the further transcription and translation of key proteins to maintain a balanced excitation/inhibition circuit. Moreover, it is important to add that this circuit can be tightly regulated by endocannabinoids [85] and by CB1R antagonists [86].

## Limitations

A few limitations must be taken into consideration while evaluating the results of this study. Firstly, we characterized at the transcriptomic level the synaptoneurosomal fractions to study synaptic RNAs. This enriched fraction could include fractions of other cell types apart from pre- and postsynaptic neuronal components. Therefore, we cannot rule out that the alternative splicing alterations observed could happen in other cell types apart from neurons. Then, further investigation will be critical to uncover whether our findings are specific of rimonabant treatment or depend on the specific CB1R inhibition at the synapses. Rimonabant is a systemic CB1R antagonist/inverse agonist that was withdrawn from the market and can only be used experimentally. However, our results were obtained with a very low dose, previously reported to alleviate FXS phenotypic traits [15]. Thus, although rimonabant at high doses could have some side effects in specific populations [87], in this study we demonstrate that this very low dose modified FX synaptic transcriptome without altering WT transcriptome. Finally, while *in silico* detections of AS changes clearly demonstrated the reversion in the direction of many AS events, we observed more subtle changes when validated in a different set of samples. It is possible that these AS events were more efficiently detected by high-throughput sequencing where millions of reads cover the event rather than with TaqMan probes, in which the number of molecules detectable are lower. It also remains unclear whether the magnitude of these AS events can vary between individuals or whether could differ between cell types. We identified changes in AS events in hippocampal synaptoneurosomes that were sensitive to rimonabant treatment, but further studies need to be done to stablish the connection between synaptic AS changes and the improvement of FXS phenotypic traits.

## Conclusions

Loss of FMRP results in changes in expression patterns in hippocampal synaptoneurosomes, which are most apparent when transcripts are independently analyzed and AS events are considered. In addition, as most of the genes that were alternatively spliced in FX sensitive to rimonabant intervention were RBPs, we propose that these genes may play an intermediate role in the translation of genes directly associated with synaptic modifications. Altogether, our results suggest that the inhibition of CB1R with rimonabant modifies mRNA splicing in the FXS synapse which could be related to the phenotypic improvement previously demonstrated by this pharmacological intervention.

## List of abbreviations

A3: Alternative 3’ splice site
A5: Alternative 5’ splice site
AF: Alternative first exon
ANOVA: Analysis of variance
AS: Alternative splicing
ASD: Autism spectrum disorder
CA1: *Cornu Ammonis* 1
CB1R: Cannabinoid type-1 receptor
CPM: Counts per million
DGE: Differential gene expression
dPSI: Delta percent spliced in
DTE: Differential transcript expression
ECS: Endocannabinoid system
FDR: False discovery rate
*FMR1*: Fragile X messenger ribonucleoprotein 1 human gene
*Fmr1*: Fragile X messenger ribonucleoprotein 1 mouse gene
FMRP: Fragile X messenger ribonucleoprotein 1
FX: *Fmr1* KO mouse
FXS: Fragile X syndrome
GO: Gene ontology
*Hnrnpd*: Heterogeneous nuclear ribonucleoprotein D mouse gene
HNRNP: Heterogeneous nuclear ribonucleoproteins
i.p.: intraperitoneally
LTD: Long-term depression
LTP: long-term potentiation
*Mff*: Mitochondrial fission factor mouse gene
MFF: Mitochondrial fission factor protein
mRNA: Messenger ribonucleic acid
MX: Mutually exclusive exons
PSI: Percent spliced in
qRT-PCR: Quantitative reverse transcriptase polymerase chain reaction
RBP: RNA binding protein
RI: Retained intron
RIM: Rimonabant
RNA-Seq: Ribonucleic acid sequencing
SE: Skipping exon
s.e.m.: Standard Error of Mean
TPM: Transcript per million
UTR: Untranslated region
VEH: Vehicle
WT: Wild-type

## Declarations

### Ethics approval and consent to participate

Animal procedures were conducted following “Animals in Research: Reporting Experiments” (ARRIVE) guidelines and standard ethical guidelines (European Directive 2010/63/EU), and were approved by the local ethical committee (Comitè Ètic d’Experimentació Animal-Parc de Recerca Biomèdica de Barcelona, CEEA-PRBB).

### Consent for publication

Not applicable

### Availability of data and materials

RNA-Seq raw-data was deposited into the Gene Expression Omnibus database under accession number GSE284627 for hippocampal synaptoneurosomes and bulk hippocampus from the same animal and GSE285032 for WT and FX hippocampal synaptoneurosomes treated with vehicle or rimonabant (0.1 mg/kg, 7 d).

## Author’s contribution

L.R.-R., *In silico* analysis, biochemical experiments, sample processing, statistical analysis, graphs and writing of the manuscript; L.C.-A., Synaptoneurosomes *in silico* characterization; A.B.-M. and L.G.-L., Biochemical experiments and sample processing; M.R.-S. and E.E., Splicing *in silico* study; S.M.-T. and A.N.-R., Animal model and sample processing; R.M., Supervision and funding of the study; A.O., Conceptualization, supervision, funding of the study and writing of the manuscript. All authors revised the final version of the manuscript.

## Acknowledgements

We are grateful to Raquel Martín, Dulce Real and Francisco Porrón for expert technical assistance and the Genomics Core Facility at Universitat Pompeu Fabra for helpful discussion.

