## Supplementary Information for "CB1 receptor inhibition in fragile X syndrome mice impacts alternative splicing alterations in hippocampal synaptoneurosomal transcriptome"

### Additional Information

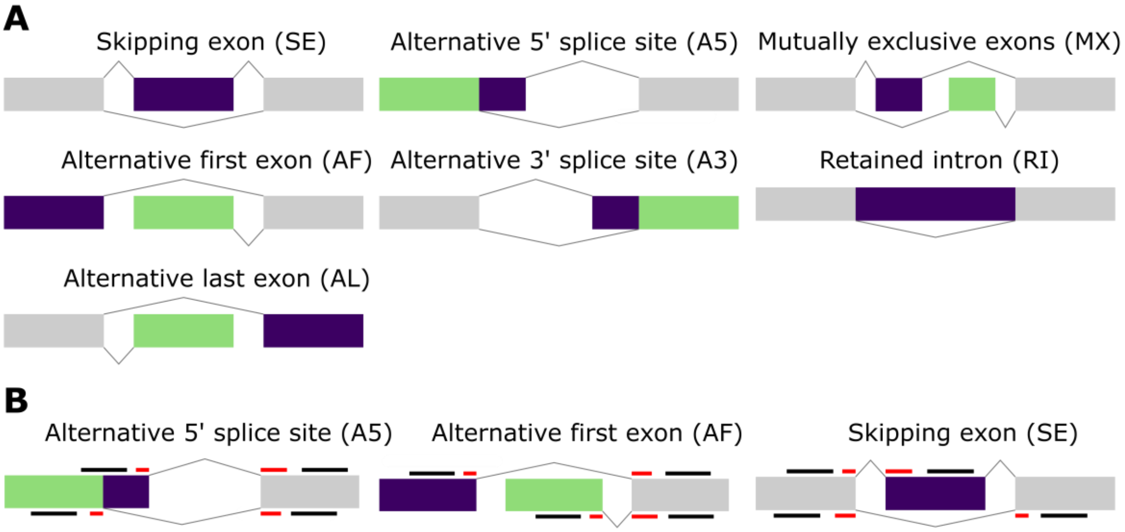


**Supplementary figure 1: *In silico* and molecular detection of AS events. (A)** Schematic representation of AS events detected by *SUPPA* with constitutive exon (grey), exon inclusion (purple) and alternative exon (green). **(B)** Schematic illustration of Taqman assays with primers (black) and internal probes (red) location for different type of event.

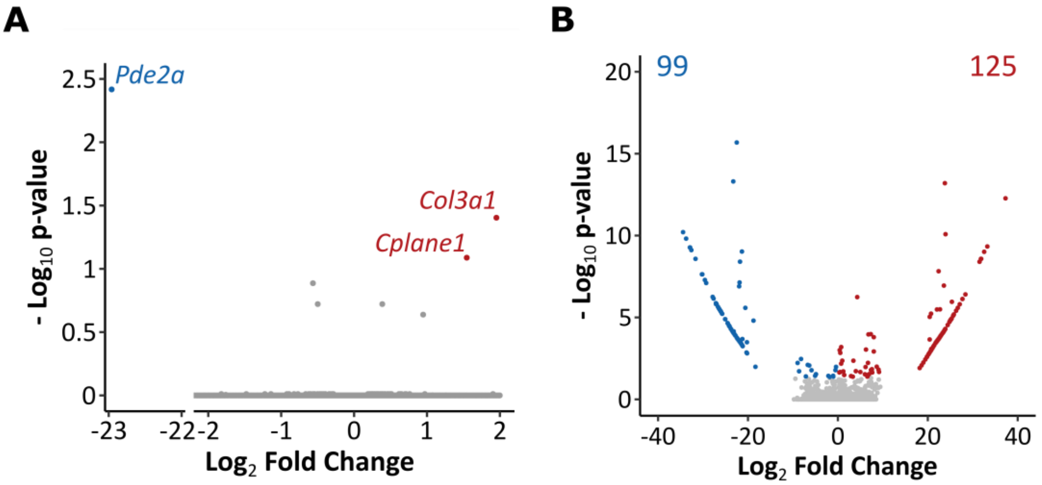


**C**

| ID | Description | Domain | P-value | Adjusted p-value | Genes |
|---|---|---|---|---|---|
| GO:0097546 | Ciliary base | CC | 1.02e-04 | 2.90e-02 | *Prkar2b, Prkacb, Ift46, Ift122* |
| GO:0047485 | Protein N-terminus binding | MF | 2.25e-04 | 4.80e-02 | *Pex5, Men1, Fez1, Tsc1, Plec, Zmynd8* |
| GO:0070064 | Proline-rich region binding | MF | 3.10e-04 | 4.50e-02 | *Sh3d19, Cyld, Abi2* |
| GO:0051018 | Protein kinase A binding | MF | 4.80e-04 | 4.60e-02 | *Prkar2b, Prkacb, Pakap, Akap12* |

**Supplementary figure 2: Transcriptomic changes in WT hippocampal synaptoneurosomes after systemic CB1R inhibition**. Volcano plot of differentially expressed **(A)** genes and **(B)** transcripts in WT synaptoneurosomes treated with rimonabant (0.1 mg/kg, 7 d) or vehicle (WT-VEH vs. WT-RIM). Red and blue dots represent significantly up or downregulated genes, respectively. (WT-VEH, n = 4; WT-RIM, n = 5). Cut-off was set at $|log2FC| > 0.2$ and adjusted *p*-value < 0.1 for genes and *p*-value < 0.05 for transcripts. **(C)** Gene ontology enrichment results from upregulated transcripts in WT synaptoneurosomes treated with rimonabant (0.1 mg/kg, 7 d). Based on GO categories: CC, cellular component; MF, molecular function.

| Type of AS | WT-VEH vs. FX-VEH | FX-VEH vs. FX-RIM | WT-VEH vs. WT-RIM |
|---|---|---|---|
| Skipping exon | 75 (69) | 47 (44) | 52 (49) |
| Alternative first exon | 85 (64) | 66 (42) | 72 (56) |
| Alternative last exon | 1 (1) | NA | NA |
| Alternative 5’ splice site | 43 (43) | 24 (23) | 32 (30) |
| Alternative 3’ splice site | 47 (44) | 31 (29) | 38 (36) |
| Mutually exclusive exons | 3 (3) | 1 (1) | 2 (2) |
| Retained intron | 5 (5) | 9 (8) | 1 (1) |

**Supplementary figure 3: AS modification in hippocampal synaptoneurosomes**. Changes in AS events and genes (in brackets) in FX and WT hippocampal synaptoneurosomes treated with vehicle (VEH) or rimonabant (0.1 mg/kg, 7 d) (RIM). Cut-off was set at |dPSI| > 0.05 and *FDR* < 0.05.
